# CRISPR/Cas9-mediated mutagenesis of phytoene desaturase in diploid and octoploid strawberry

**DOI:** 10.1101/471680

**Authors:** Fiona Wilson, Kate Harrison, Andrew D. Armitage, Andrew J. Simkin, Richard J. Harrison

**Affiliations:** NIAB EMR, New Road, East Malling, Kent, ME19 6BJ, UK

**Keywords:** *Fragaria*, strawberry, diploid, polyploid, CRISPR/Cas9, PDS

## Abstract

Gene editing using CRISPR/Cas9 is a simple and powerful tool for elucidating genetic controls and for crop improvement. We demonstrate use of CRISPR/Cas methodology in diploid *Fragaria vesca* ssp. *vesca* ‘Hawaii 4’ and octoploid *F. x ananassa* ‘Calypso’ by targeting the visible endogenous marker gene *PDS* (phytoene desaturase). *Agrobacterium*-mediated transformation of leaf and petiole explants was used for efficient stable integration of constructs expressing plant codon-optimised Cas9 and single guide sequences under control of either the *Arabidopsis U6-26* consensus promoter and terminator or *Fragaria vesca U6III* regulatory sequences. More than 80% (‘Hawaii 4’) and 50% (‘Calypso’) putative transgenic shoot lines exhibited mutant phenotypes. Of mutant shoot lines selected for molecular analysis, approximately 55% (‘Calypso’) and 75% (‘Hawaii 4’) included albino regenerants with bi-allelic target sequence variants. Our results indicate the *PDS* gene is functionally diploid in ‘Calypso’ and clearly demonstrate that CRISPR/Cas9 can be used to edit single copy genes at high frequency within the genome of the diploid and the same target in octoploid strawberry.

## 1. Introduction

The cultivated strawberry, *Fragaria* x *ananassa*, is one of the most important fruit crops in the family Rosaceae. It is among the most widely grown and consumed fruit throughout the world, with global production reaching 9.1M tonnes in 2016, an increase over the previous decade of more than 5% annually (www.freshfruitportal.com). In the UK alone, in 2015 115,000 tonnes were produced, with a market value of £253 million [1].

Traditional breeding is lengthy and difficult as *F*. x *ananassa* is an octoploid species with a complex genome, and is intolerant to inbreeding. Important for genetic improvement is the potential to enhance elite cultivars using genetic modification, first demonstrated with marker genes using *Agrobacterium*-mediated transformation in the 1990s [2, 3]. Since then a number of developments in transformation methodology and its application in *Fragaria* have been reported (see [4, 5]. The diploid wild strawberry *F. vesca* shares a high degree of sequence identity with the cultivated strawberry and is a model for genetic improvement in the genus due to its small genome size (240 mb), short generation time, and ease of genetic transformation [6, 7]. The availability of the complete genome for *F. vesca* [8, 9] and partial genome sequence of the octoploid genome [10] facilitates the identification and manipulation of genes controlling important traits.

From its first use in model species such as *Arabidopsis* [11, 12] and tobacco [12, 13], the rapid development of CRISPR/Cas9 technology has enabled widespread use of precise gene editing in plants [14-16]. Reports of successful gene editing in diploid crops include rice [17, 18], sorghum [19], tomato [20], maize [21, 22], *Populus* [23], apple [24], grapes [25] and cassava [26, 27]. In polyploid crops, multiple gene homoeoalleles have been simultaneously edited using CRISPR/Cas9 in wheat [28], potato [29] and *Brassica napus* [30, 31]. Multiple gene targets have been successfully targeted in Arabidopsis [32], maize [33] and rice [34, 35].

Defects in *PDS* gene function result in a distinctive albino phenotype [36], and recent studies in *Populus* [23], apple [24], grapes [25] and cassava [26] have targeted the *PDS* gene to demonstrate successful genome editing in these crops. Nishitani *et al*. [24] compared target sites in different exons of the apple *PDS* gene and showed that the greatest mutation efficiency was achieved by targeting exon 7. As strawberry is also a member of the family *Rosaceae*, we chose to target the same region in the *PDS* gene of diploid and octoploid strawberry.

Our results show the mutation of PDS in diploid and octoploid strawberry results in a clear albino phenotype at a high frequency. The data provided will inform future work in *Fragaria* using CRISPR/Cas9, both as a research tool and for modifying molecular mechanisms controlling traits of agronomic importance, where mutant phenotypes cannot be identified visually. The importance of this technology for use in creating new varieties in support of global food security has been recently reviewed [37].

## 2. Materials and Methods

### 2.1 Construction of CRISPR/Cas9 vectors

The binary vector pFGC-pcoCas9 (Addgene plasmid # 52256) was modified by insertion of pGreen plant marker constructs [38]: the ‘Nos-Kan’ cassette (comprising the *nptII* gene under control of nopaline synthase regulatory sequences) was cloned into the *Pme*1 site near the right border and the ‘35S-GFP’ cassette (comprising the modified *GFP* gene, [39], under control of *35S* regulatory sequences) was cloned into the *Sma*1 site in the multiple cloning site to give pCas9-K-GFP (Fig. 1).

**Fig. 1.**
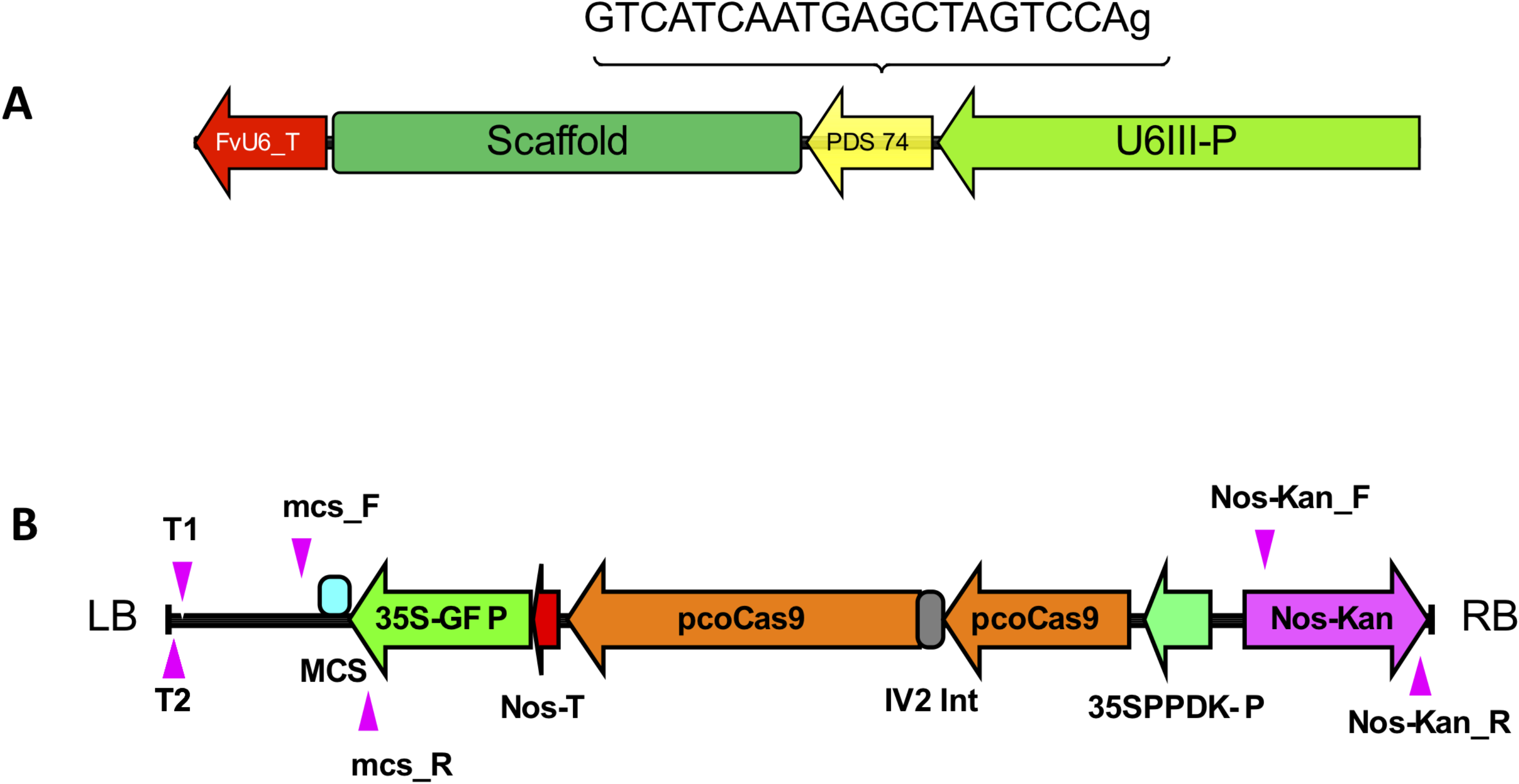
Schematic map of CRISPR cassette and T-DNA of pCas9-K-GFP. (A) Single guide RNA inserted into the multiple cloning site (MCS) of pCas9-K-GFP. (B) pCas9-K-GFP T-DNA showing the position of marker/selection cassettes and pcoCas9 coding sequence between the left border (LB) and right border (RB), and the binding regions of primers used for TAIL-PCR (T1=Tail 1; T2=Tail 2), sgRNA insertion site PCR (mcs_F; mcs_R) and Nos-Kan cassette PCR (Nos-Kan_F; Nos-Kan_R).

CRISPR cassettes incorporating the sgRNA sequence and regulatory elements (Fig. 1) were synthesised by Integrated DNA Technologies (IDT) Ltd and cloned into pCR^®^-Blunt II-TOPO^®^ (Thermo Fisher Scientific, UK). Cassettes were inserted into the *Eco*R1 site of pCas9-K-GFP and the orientation confirmed by sequencing. The *PDS* coding sequence for *Malus domestica* (GenBank: KU508828.1) was BLAST aligned against the genomes for *F. vesca* (FvAssembly 1.1) and *F*. x *ananassa* (www.rosaceae.org). The CRISPR target sequence selected (ACCTGATCGAGTAACTACTG<u>AGG</u>) is common to both *Fragaria* genomes and is located on the sense strand in exon 7 of FvAssembly 4 [9] (Fig. 2). It corresponds almost exactly to the target sequence (ex7-20bp, ACCTGATCGAGTAACTACAG<u>AGG</u>) used successfully by Nishitani *et al*. [24]. The PAM sequences are underlined. The target guide RNA (gRNA) sequence (PDS74) comprises the first 20 base pairs of the target sequence preceded by an additional guanine base to maintain the native *U6* promoter start of transcription. This is placed under transcriptional control of the consensus sequence of the *Arabidopsis U6-26* promoter [13] or the corresponding sequence of the *F. vesca U6III* promoter and terminator sequences (Table S1).

**Fig. 2.**
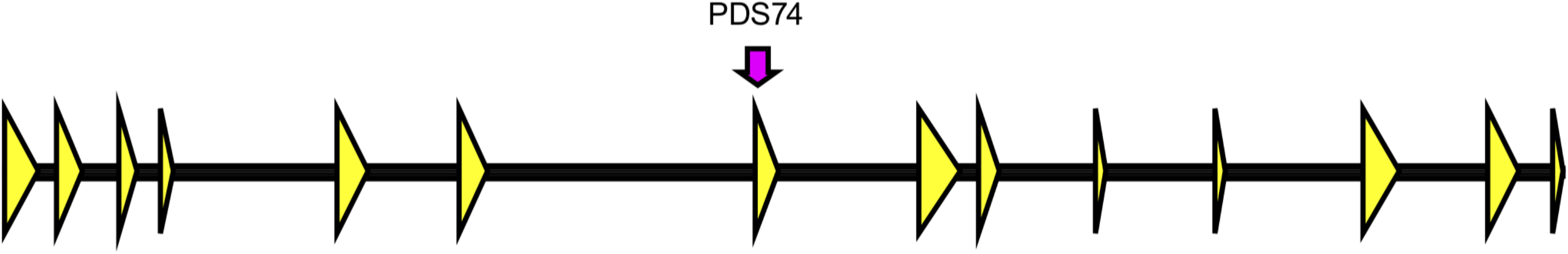
Schematic map of *Fragaria vesca PDS* gene. LG4-gene12690 (8,035 bp, FvAssembly v4) showing exons (yellow arrows) and the location of the CRISPR target sequence PDS74 (position 3’-5’ = 3,873-3,892).

### 2.2 In vitro culture of Fragaria stock cultures

*In vitro* shoot cultures of *F. vesca* ‘Hawaii 4’ and *F. x annanassa* ‘Calypso’ were maintained in a growth room at 21 ^o^C with a 16/8 h light/dark photoperiod provided by fluorescent lamps (colour reference 835, colour temperature 3500K). Crowns were subcultured at 4-5 week intervals, 5 per honey jar containing 50 ml medium. Basal culture medium was Murashige and Skoog (MS) macro and micro elements and vitamins [40], supplemented with sucrose (30 g l^−1^), solidified with Daishin agar (Duchefa D1004, 9 g l^−1^). The pH was adjusted to 5.8 before autoclaving. Shoots were alternately cultured on basal −1 medium supplemented with 6-benzylaminopurine (BAP) 0.1 mg l^−1^ and indole-3-butyric acid (IBA) 0.1 mg l^−1^ or BAP 0.5 mg l^−1^, as described by Schaart [41].

### 2.3 Growth and transformation of diploid and octoploid strawberry

Transformation of ‘Calypso’ was performed essentially as Schaart [44], with minor modifications. *Agrobacterium tumefaciens* strain EHA105 [42] harbouring the binary vector was grown overnight, and pelleted at 2,000 xg for 10 minutes. The inoculum was prepared by re-suspending the overnight culture in filter-sterilised liquid MS-based medium supplemented with glucose (30 g l^−1^) and acetosyringone (100 μM), pH 5.2, to give OD 600 nm 0.2 – 0.3. Petioles from apical leaves of *F. vesca* ‘Hawaii 4’ or leaflets from young expanding leaves of *F*. x *ananassa* ‘Calypso’ were harvested from shoots four weeks after subculture and submerged in the inoculum for approximately 10-15 minutes during explant preparation: petioles were cut into 4-5 mm pieces and leaflets were separated from each leaf before scoring transversely into 2 mm strips, leaving one leaf edge intact. Explants were then blotted to remove excess inoculum before transfer to Shoot Regeneration Medium (SRM): MS medium supplemented with α-naphthaleneacetic acid (NAA) (0.2 mg l^−1^) and thidiazuron (TDZ) (1 mg l^−1^). The pH was adjusted to 5.8 and Agargel^™^ (A3301, Sigma) (5 g l^−1^) was added before autoclaving. After autoclaving filter-sterilised glucose (300 g l^−1^) was added to a final concentration of 30 g l^−1^. After four days’ dark incubation at 21 ^o^C, explants were washed in a solution of filter-sterilised ticarcillin disodium/clavulanate potassium (TCA, Duchefa T0190) (400 mg l^−1^) in water, blotted and transferred to SRM selection medium containing filter-sterilised TCA (400 mg l^−1^) and kanamycin sulphate monohydrate (Kanamycin, Duchefa K0126) (100 mg l^−1^). ‘Hawaii 4’ petioles were initially cultured in Sarstedt Cell Culture Flasks (T-25) containing 15 ml liquid selection medium, for four weeks, 30 - 50 explants per flask, shaking at 60 rpm at low light intensity, before transfer to semi-solid selection medium. Leaves were cultured throughout on semi-solid selection medium: approximately 10 leaves (abaxial side in contact with the medium) or 20 petiole pieces were cultured per 90mm triple-vent Petri dish containing approximately 30 ml medium. Dishes were sealed with Parafilm_®_ M and incubated in the growth room as for shoot cultures. Subculture intervals were 4-6 weeks. At the second culture passage, half of the explants were transferred to SRM supplemented with gibberellic acid (GA_3_) (1 mg l^−1^) before autoclaving. Petioles were divided to separate calli developing at both petiole ends. Leaves were divided as expansion occurred to ensure contact with the selection medium, to facilitate subculturing and to separate regenerating calli. Calli with shoots were harvested over a 7-month period, and transferred to rooting medium: MS medium 2.2 g l^−1^, supplemented with sucrose 20 g l^−1^, BAP 0.1 mg l^−1^, IBA 0.1 mg l^−1^, solidified with 9 g l^−1^ Daishin agar, pH adjusted to 5.8 before autoclaving. Filter-sterilised kanamycin (100 mg l^−1^) and TCA (400 mg l^−1^) were added after autoclaving. For shoots with an obvious albino phenotype, kanamycin was omitted from the medium. Albino and variegated shoots were subcultured onto rooting medium with TCA (400 mg l^−1^).

### 2.4 Analysis of transgenic lines

DNA was extracted from leaf material using the DNeasy Plant Kit (Qiagen, Manchester, UK). Primers used in PCR and sequencing reactions (Table S2) were synthesised by IDT Ltd. PCR screening of putative transgenic lines for Nos-Kan cassette and sgRNA insertions was performed using PCRBIO Taq Mix Red (PCR Biosystems) and reaction conditions as detailed in Table S3. PCR amplicons spanning the target region or Left Border T-DNA-genomic DNA junction sites were sequenced by Illumina MiSeq: primers incorporated target-specific sequences at the 3’ ends, and Illumina adapter overhang sequences (P5 forward overhang, P7 reverse overhang) at the 5’ ends. A 401 bp genomic region was amplified using primer pair P5-300 F and P7-700F, binding 186 - >161 bp upstream and 195 -> 170 downstream of the target region, respectively. Amplicons were prepared for sequencing following Illumina guide 16S Metagenomic Sequencing Library Preparation (Part # 15044223 Rev. B). Template was amplified using Q5^®^ Hot Start High-Fidelity 2X Master Mix (M0494, New England Biolabs Ltd.) and purified using Mag-Bind^®^ TotalPure NGS (M1378, Omega Bio-tek Inc.). Dual indices and Illumina sequencing adapters were attached using Nextera XT Index Kit (15055290, Illumina) and 2x KAPA HiFi HotStart ReadyMix (KK2605, Kapa Biosystems). PhiX control V3 (15017660, Illumina) and the amplicon library were diluted to 4 pM concentration and the sample was spiked with 30% PhiX before loading onto a MiSeq Reagent Nano Kit v2 500 cartridge (MS-103-1003, Illumina).

Thermal Asymmetric Interlaced (TAIL) PCR was used to amplify T-DNA-genomic DNA junctions using PCRBIO Taq Mix Red (PCR Biosystems Ltd): first round PCR was performed using arbitrary degenerate primer AD3 [43] and T-DNA-specific primer TAIL R1 (binding 257 -> 228 downstream of the Left Border). Nested primer TAIL R2 (binding 209 -> 188 downstream of the left border) and AD3 were used for the second round PCR. PCR reactions were performed using a Verity 96-Well Thermal Cycler (#4375786, maximum block ramp of 3.9 °C/Sec and a maximum sample ramp of 3.35 °C/Sec). The PCR cycle was essentially as Liu et al., 1995, with minor modifications (Table S3). Amplicons were purified using Mag-Bind^®^ TotalPure NGS and Sanger-sequenced by Eurofins Genetic Services Ltd. The sequencing primer (TAIL SEQ) binds 177 -> 146 downstream of the T-DNA left border.

### 2.5 Sequencing analysis and software

Illumina sequencing reads were trimmed to remove low quality data and Illumina adapters with fastq-mcf (http://code.google.com/p/ea-utils) before being aligned to the reference *F. vesca* ‘Hawaii 4’ genome v4.0.a1 [9], using BWA v0.7.15 [45]. Variants in respect to the reference genome were predicted and quantified from aligned reads using the CrispRVariant package v1.9.2 [46]. Nucleotide and protein alignments were performed using Geneious version 10.0.2 (http://www.geneious.com). [47]. Schematic maps were prepared using IBS software [48].

## 3. Results and Discussion

### 3.1 Transformation and regeneration of mutant lines

We placed sgRNAs under transcriptional control of the Arabidopsis *U6-26* promoter (p*AtU6-26* vector) [13] or the corresponding sequence of the *F. vesca U6III* promoter (p*FvU6III* vector).

Following transformation, Calli, which produced shoots were maintained on selective regeneration medium for up to seven months and shoots were successively harvested as they regenerated. Calli were regenerated on media with or without GA_3_ to encourage rooting. Mutant shoots were efficiently obtained on medium without GA_3_ and there is no clear indication of the effect of including GA_3_ in the regeneration medium (Table 1). Up to 90% of ‘Hawaii 4’ and ‘Calypso’ shoot lines (shoots from isolated calli) formed roots on kanamycin or exhibited clear mutant phenotypes (Table 1 and Fig. 3, 4).

**Table 1.** Numbers and percentages of lines that regenerated in *Fragaria vesca* and *Fragaria x ananassa* depending on construct promoters and GA treatment

| Promoter | Regeneration medium +/- GA<sub>3</sub> | Number of lines harvested throughout the experiment |  |  |  | Total Lines harvested | Number of lines assessed for transformed phenotype | Putative transformed lines | % putative transformed lines | Lines with mutant phenotype | % putative transformants with mutant phenotype | Cultivar |
| --- | --- | --- | --- | --- | --- | --- | --- | --- | --- | --- | --- | --- |
|  |  | 2 months | 4 months | 5 months | 7 months |  |  |  |  |  |  |  |
| At | + GA | 18 | 18 | 22 | 7 | 65 | 58 | 42 | 72 | 24 | 57 | Hawaii-4 |
| At | no GA | 1 | 4 | 1 | 0 | 6 | 6 | 5 | 83 | 1 | 20 |  |
| Fv | + GA | 12 | 1 | 39 | 4 | 56 | 55 | 50 | 91 | 46 | 92 |  |
| Fv | no GA | 15 | 8 | 15 | 0 | 38 | 37 | 26 | 70 | 21 | 81 |  |
| Average |  |  |  |  |  |  |  |  | 79 |  | 62 |  |
| At | + GA | 0 | 11 | 10 | 21 | 42 | 30 | 22 | 73 | 8 | 36 | Calypso |
| At | no GA | 0 | 9 | 7 | 6 | 22 | 19 | 16 | 84 | 11 | 69 |  |
| Fv | + GA | 2 | 4 | 1 | 6 | 13 | 9 | 8 | 89 | 4 | 50 |  |
| Fv | no GA | 2 | 3 | 4 | 3 | 12 | 10 | 8 | 80 | 2 | 25 |  |
| Average |  |  |  |  |  |  |  |  | 82 |  | 45 |  |

**Fig. 3.**
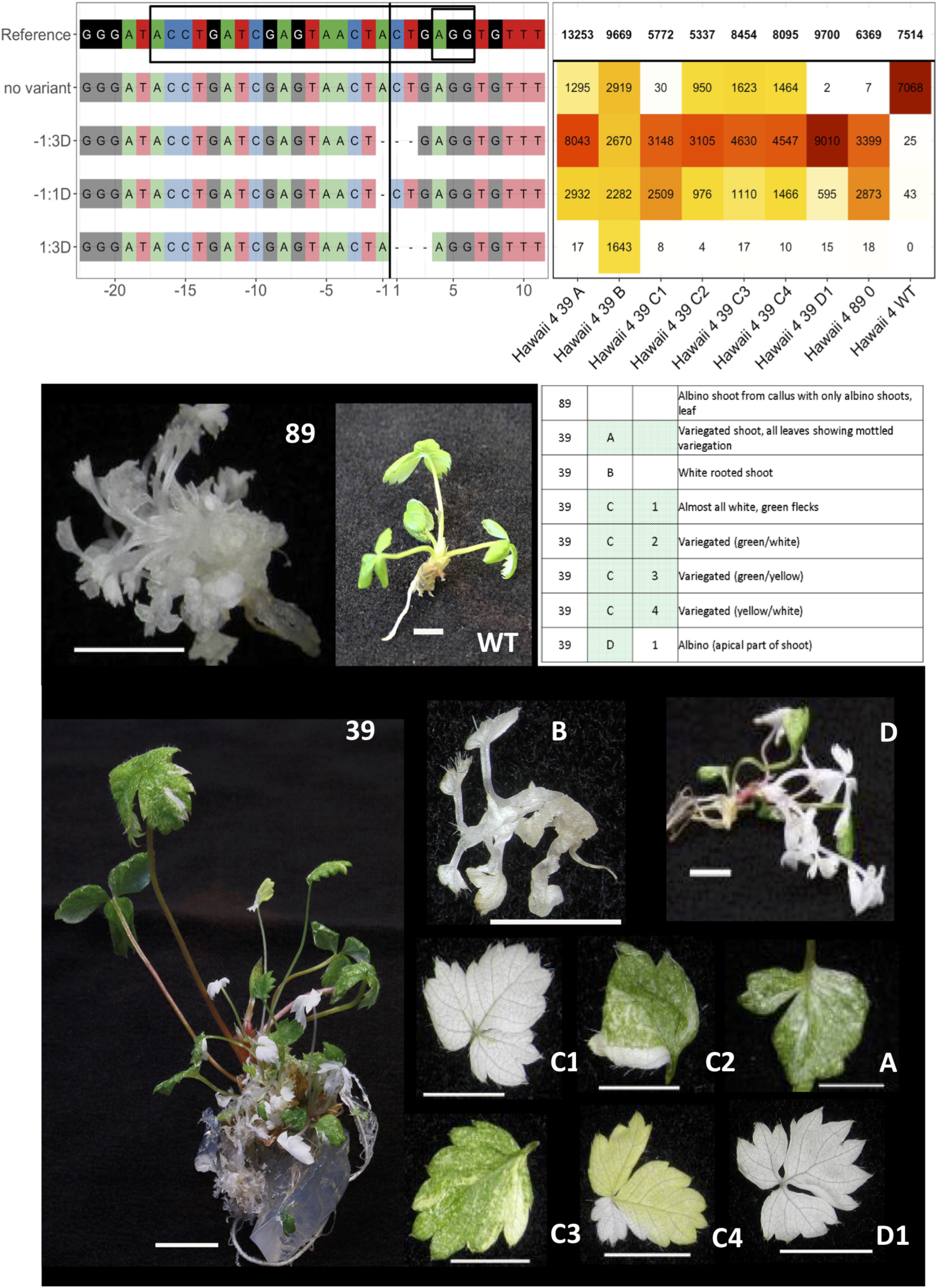
Sequence mutations and corresponding phenotypes of CRISPR/Cas9 transgenic lines of ‘Hawaii 4’. Examples of sequence mutations and allelic variants with corresponding phenotypes in transgenic shoot lines of *Fragaria* ‘Hawaii 4’ (line numbers 39 and 89). Each sequence variant type is identified by the location of the mutation relative to the cut site:number bases deleted (D). Deletions result in amino acid loss (mutations −1:3D and 1:3D) and also frameshift (mutation −1:1D) and generation of stop codons within exon 7, resulting in truncated PDS protein. The cut site is shown by a black vertical line; the target site and PAM site (AGG) are within the box in the reference sequence. WT=wild type. Scale bars are 5 mm.

**Fig. 4.**
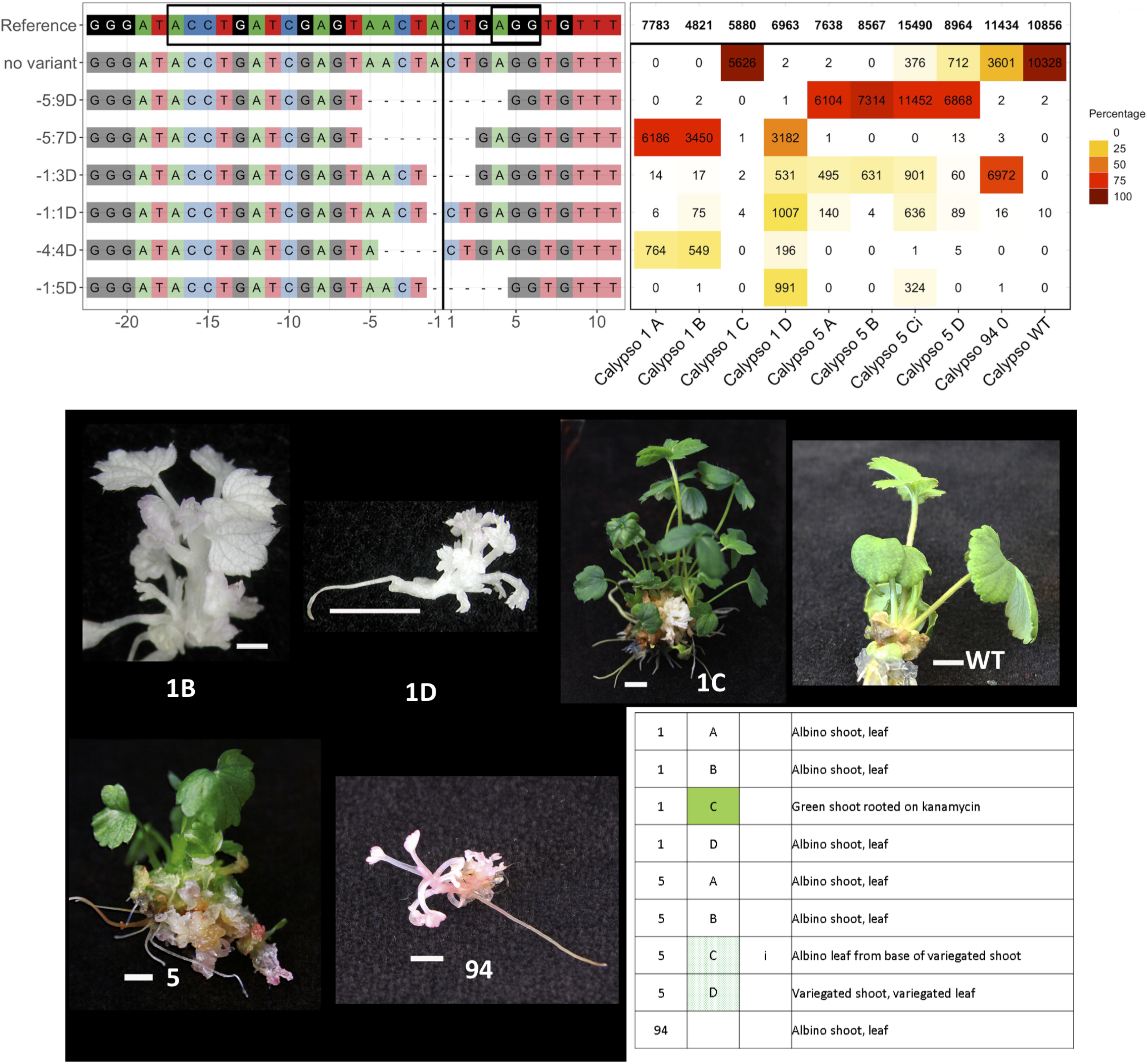
Sequence mutations and corresponding phenotypes of CRISPR/Cas9 transgenic lines of ‘Calypso’. Examples of sequence mutations and allelic variants with corresponding phenotypes in transgenic shoot lines of *Fragaria* ‘Hawaii 4’ (line numbers 39 and 89). Each sequence variant type is identified by the location of the mutation relative to the cut site:number bases deleted (D). Deletions result in amino acid loss (mutations −1:3D and −5:9D) and also frameshift (mutation −1:1D, −1:5D, −4:4D and −5:7D) and generation of stop codons within exon 7, resulting in truncated PDS protein. The cut site is shown by a black vertical line; the target site and PAM site (AGG) are within the box in the reference sequence. WT=wild type. Scale bars are 5 mm.

Approximately 62% (‘Hawaii 4’ Fig 3) and 45% (‘Calypso’ Fig. 4) putative transgenic shoot lines exhibited mutant phenotypes (Table 1). Where shoot lines comprised a range of phenotypes, albino or variegated phenotypes were apparent in approximately 10% of tissues harvested. After callus formation, half of explants were placed on regeneration medium supplemented with GA3, as it was unknown whether mutation in the *PDS* gene would inhibit gibberellic acid synthesis and adversely affect shoot development. Mutant shoots were efficiently obtained on medium without GA3 and there is no clear indication of the effect of including GA3 in the regeneration medium. We placed sgRNAs under transcriptional control of the *Arabidopsis U6-26* promoter (p*AtU6-26* vector) [13] or the corresponding sequence of the *F. vesca U6III* promoter (p*FvU6III* vector). The proportions of ‘Hawaii 4’ and ‘Calypso’ mutant phenotypes derived from p*AtU6-26* are 25/47 lines (53%) and 19/38 lines (50%), respectively, and corresponding data for p*FvU6III* are 67/76 lines (88%) and 6/16 lines (38%). Native promoters are known to enhance mutation efficiency: for example, in soybean in three target genes, mutation efficiencies were increased by the use of the soybean U6-10 promoter compared to the Arabidopsis U6-26 promoter [49]. Relative mutation efficiencies in this experiment suggest that the *FvU6III* promoter may be more effective than the *AtU6-26* consensus promoter in ‘Hawaii 4’ but not in ‘Calypso’, and that alternative U6 promoters native to the octoploid genome may enhance sgRNA expression in ‘Calypso’ and other commercial cultivars.

### 3.2 Analysis of transgenic lines

#### 3.2.1 Types and frequency of target site mutations

Some 401 bp amplicons were generated for 96 samples, including wild-type ‘Hawaii 4’ and ‘Calypso’ (Fig. S1). Illumina MiSeq results were obtained for both wild types, 18 ‘Hawaii 4’ and 7 ‘Calypso’ transgenic shoot lines. Transgenic shoot lines exhibit a variety of mutant sequence variants at and around the target site in a highly conserved region of the *PDS* gene (Figs. 3, 4, S2, S3), resulting in phenotypes typical of defective *PDS* gene function, including a high proportion of completely albino phenotypes. Approximately 60-80% shoot lines regenerated albino shoots with 100% target site sequence variant reads.

The most common sequence mutations in ‘Hawaii 4’ (H4) and ‘Calypso’ are three variant types with 1, 2 or 3 base deletions at (or spanning) the cut site in exon 7 of the *PDS* gene (Table 2, Fig. 5): all are present in the data for the majority of ‘Hawaii 4’ lines and approximately half of ‘Calypso’ lines (Fig. 6A). Insertions and substitutions are uncommon and seen only in ‘Hawaii 4’: 3 of 18 lines have a 1-base insertion and one line has a 2-base insertion; and 2 lines have single A/C substitutions at or near the cut site: line 32 has both a single base substitution and a 1-base insertion; and line 46 has just a single base substitution. Large deletions (10 bases or more) are also uncommon, present in 4 of ‘Hawaii 4’ lines. Protein translation of in-frame mutations (deletions of 3, or multiples of 3, bases) shows loss of one or more amino acids around the cut site. Frameshift mutations (and mutation −44:45D) result in amino acid loss and generation of stop codons within exon 7, resulting in truncated PDS protein.

**Table 2.** Classification of mutants within regenerated lines (presence and absence)

| Sequence variant | Hawaii 4 | Calypso | Non-synonymous<br>amino acid<br>substitution | Frameshift & protein<br>truncation | Amino acid deletion | Putative functional<br>protein |
| --- | --- | --- | --- | --- | --- | --- |
| -1:1D | 17 | 4 | - | + | + | - |
| -2:1D | 2 | - | - | + | + | - |
| -2:2D | 10 | 3 | - | + | + | - |
| -1:3D | 11 | 3 | - | - | + | - |
| -4:4D | - | 1 | - | + | + | - |
| -1:5D | 1 | 3 | - | + | + | - |
| -2:6D | 1 | - | - | - | + | - |
| -3:7D | - | 1 | - | + | + | - |
| -5:7D | - | 1 | - | + | + | - |
| -2:9D | 1 | - | - | - | + | - |
| -5:9D | 1 | 1 | - | - | + | - |
| -6:9D | 1 | - | - | - | + | - |
| -10:9D | 1 | - | - | - | + | - |
| -11:15D | 1 | - | - | - | + | - |
| -1:22D | 1 | - | - | + | + | - |
| -25:271D | 1 | - | - | - | + | - |
| -44:45D | 1 | - | - | + | + | - |
| 1:3D | 1 | - | - | - | + | - |
| 1:5D | 1 | - | - | + | + | - |
| 1:1I | 3 | - | + | + | - | - |
| SNV:-3A<br>& 1:2 | 1 | - | + | + | - | - |
| SNV:1A | 1 | - | + | - | - | + |

**Fig. 5.**
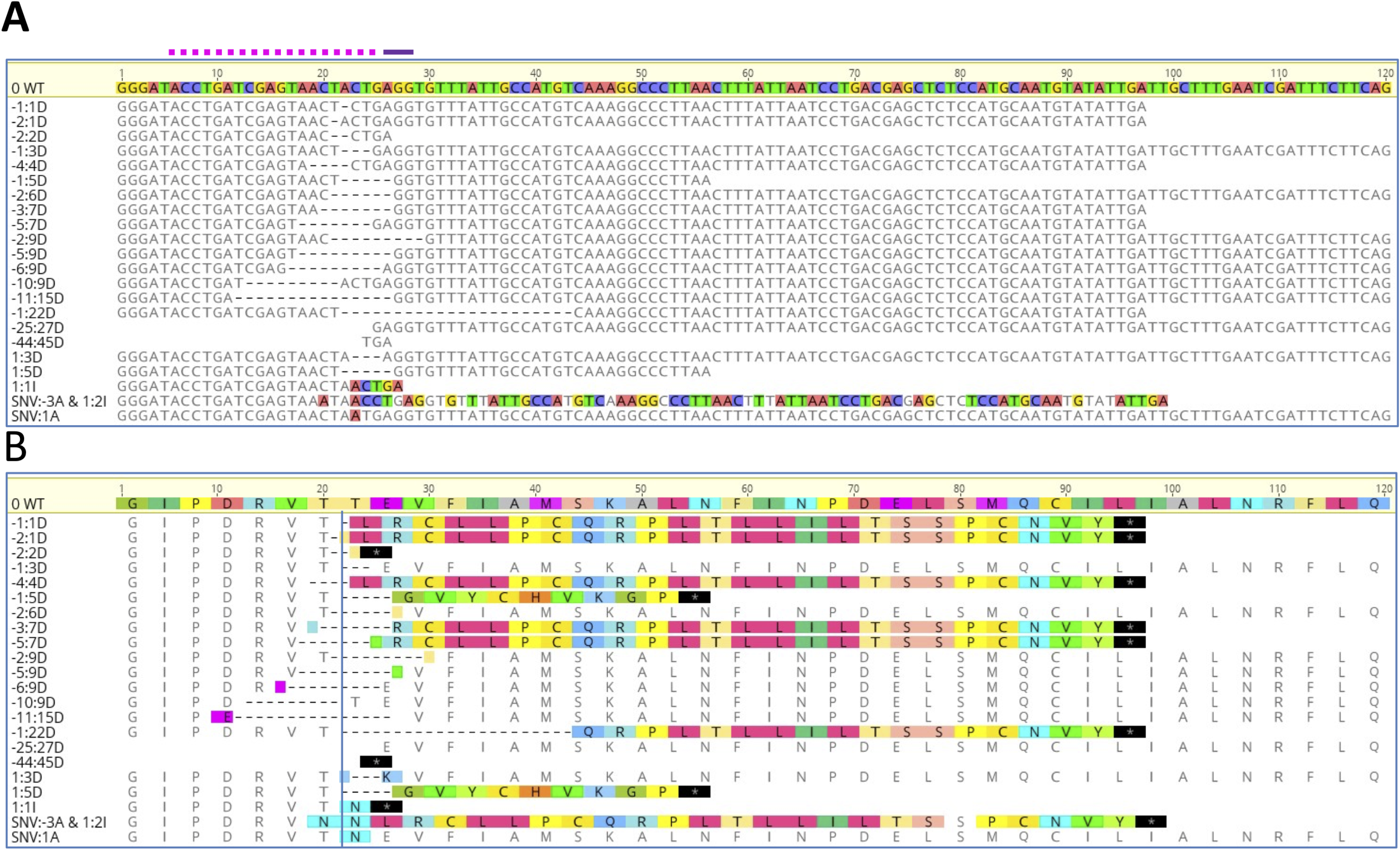
Mutations of the *PDS* gene in CRISPR/Cas9 transgenic lines of ‘Hawaii 4’ and ‘Calypso’. (A) Summary of CRIPSR/Cas9 sequence variants the *Fragaria PDS* gene, aligned to the reference (wild-type) sequence for exon 7 (120 base pairs). The wild-type (WT) nucleotide sequence is shown at the top of the chart and the positions of the 20 base pair target region ‘PDS74’ and PAM site (AGG) are indicated above the chart by dotted and solid lines, respectively. The ‘cut site’ is 3 base pairs upstream of the PAM site (AGG) and is shown as a vertical line. Each sequence variant type is identified by the location of the mutation relative to the cut site:number of bases deleted (D), inserted (I), or substituted (SNV:location relative to cut site and base substitution). Deleted bases generating gaps within exon 7 are shown by dashes. Base insertions and substitutions and consequent misaligned sequences relative to the wild type are highlighted. Sequences are shown truncated where mutations cause a frameshift generating stop codons within exon 7. (B) Amino acid sequences for each variant type aligned to the reference sequence (WT) for exon 7. Variant amino acids relative to the wild type are highlighted. Stop codons are shown in black and marked with an asterisk. Gaps in amino acid sequences within exon 7 are shown by a dashed line and partial amino acids are highlighted.

**Fig. 6.**
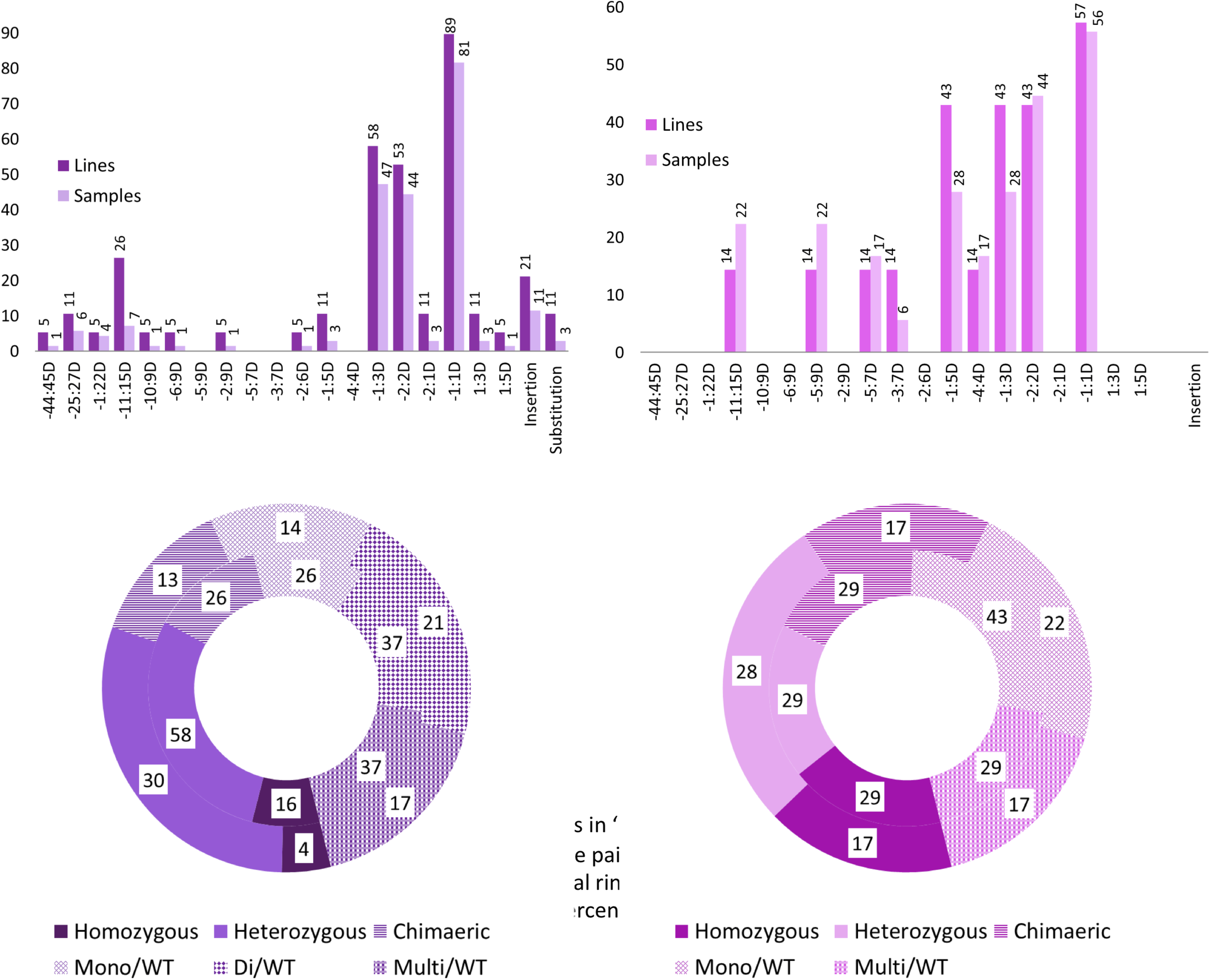
Sequence mutations and allelic variants in ‘Hawaii 4’ and ‘Calypso’ CRISPR/Cas9 transformants. (A) Sequence mutations: location of deletions relative to the cut site:number base pairs deleted, insertions (1 or 2 base pairs) and single base pair substitutions. (B) Types of allelic variant (central ring = shoot lines, outer ring = individual samples). Frequencies of sequence mutations and types of allelic variant are shown as percentages of total shoot lines and samples sequenced. Numbers of shoot lines/samples sequenced for ‘Hawaii 4’ and ‘Calypso’ are 18/69 and 7/19, respectively.

#### 3.2.2 Types of allelic variant

Samples are either homozygous for one sequence variant type, heterozygous (having two sequence variant types), chimaeric (3 or more sequence variant types), or have a mixture of wild-type sequence and 1, 2 or more than 2 sequence variants: mono-WT, di-WT or multi-WT (Table S4, Fig. 6B). All four homozygous variants are of the most common variant type (−1:1D); and 34 of 35 heterozygous allelic variants are combinations of or include the most common 1-, 2- or 3-base sequence variants (−1:1D, −2:2D, −1:2D). Most homozygous and heterozygous allelic variants are of the most common sequence variant types and are likely to have arisen from simultaneous mutation of both alleles prior to cell division. Chimaeric allelic variants indicate that mutation occurred in one allele either before cell division or at a later stage, followed by additional mutation events. Sequence data for multiple samples from single shoot lines share variant sequence types, and combinations of sequence types (Table S4, Fig. 3, 4, S2, S3). The timing and sequence of mutations will determine the mixture of the allelic variants within a shoot and shoot line.

Four of 7 ‘Calypso’ shoot lines (lines 1, 5, 7, 100) and 14 of 18 ‘Hawaii 4’ shoot lines (all lines except 9, 10, 32, 36) produced shoots which are biallelic (homozygous, heterozygous or chimaeric) mutants.

#### 3.2.3 The evidence suggests that multiple shoots from individual callus lines will often be derived from the same transgenic event

For selected samples, sequences for left border integration sites were generated by TAIL PCR (Fig. S4). Putative plant chromosomal sequence (i.e. not matching vector sequence) at the integration site, often contiguous with partial or complete T-DNA left border sequence, was recovered from approximately 40% of shoot lines screened (7 of 17 ‘Hawaii 4’ and 4 of 9 ‘Calypso’ lines, Fig. 7A, Table S5). The T-DNA integration sequences are different for each shoot line (Table S4). For approximately 50% of shoot lines sequenced (9 ‘Hawaii 4’ and 4 ‘Calypso’ shoot lines) the sequence obtained beyond the T-DNA left border matches only binary vector sequence (Fig. 7B), indicating chromosomal integration of vector ‘backbone’ DNA, which was shown by De Buck *et al*. [49] to be a common occurrence in transgenic *Arabidopsis* and tobacco plants. Shoots of ‘Hawaii 4’ line 10 have a complex integration sequence comprising T-DNA, vector backbone sequence and sequence not matching vector DNA (Fig. S5).

**Fig. 7.**
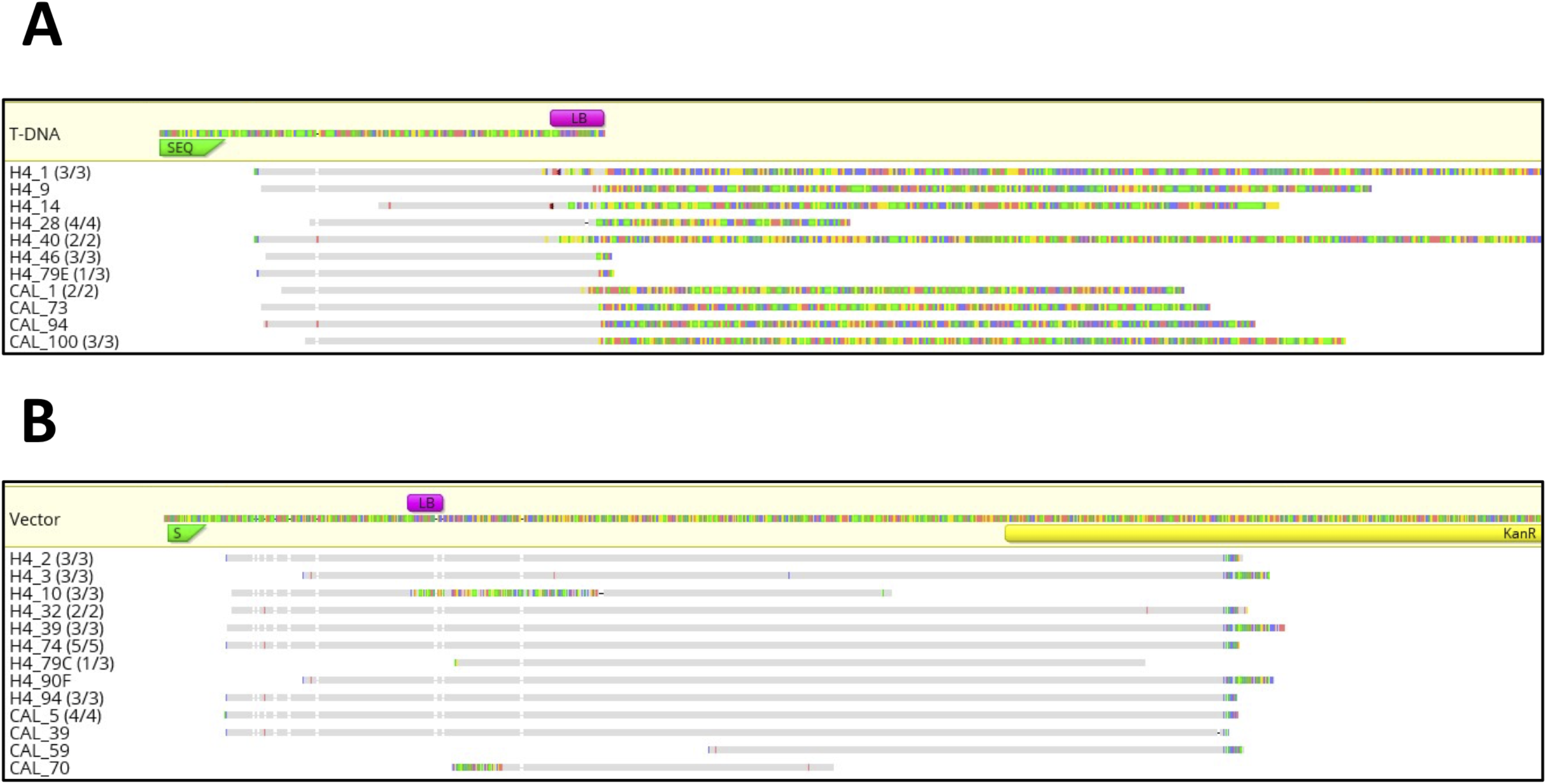
Left border T-DNA integration. TAIL PCR sequence data for transgenic ‘Hawaii 4’ (H4) and ‘Calypso’ (Cal) lines. (A) Assembly to the binary vector T-DNA. (B) Assembly to the binary vector pCas9-K-GFP. Where homologous data was obtained from more than one sample in shoot lines this is indicated (number of homologous shoots/number of shoots sequenced in shoot line) and sequence for only one (representative) sample is shown. Bases not matching the reference sequence (putative chromosomal sequence) are highlighted. S/SEQ=primer used for sequencing; LB=T-DNA left border.

The sequence data for multiple shoots from single lines (4 of 9 ‘Calypso’ lines and 13 of 17 ‘Hawaii 4’ lines) have either high homology to each other where putative genome integration sequences were obtained, or are all homologous to vector ‘backbone’ sequence (Fig. 7). The only line where shoots appear not to share a transgenic origin, is ‘Hawaii 4’ line 79: 79 C and 79 E TAIL sequences do not overlap but both align to the vector backbone (Fig. S5); 79E has a more complex pattern of mutation and does not align to vector sequence.

The data from both target site sequence analysis and TAIL PCR sequence analysis suggest that multiple shoots regenerated from individual callus lines are likely to share a transgenic integration origin.

#### 3.2.4 Observed phenotype is frequently consistent with mutation sequence data obtained

Phenotypic observations of leaf samples of multiple shoots were compared to the mutant sequence data generated for each sample (7 of 10 ‘Calypso’ and all 18 ‘Hawaii 4’ transgenic shoot lines (Figs. 3, 4, S2, S3, Table S4). All shoot lines comprised a mixture of green, and/or variegated, pale green, white or albino shoots, except ‘Hawaii 4’ line 89, for which only albino callus and shoots were regenerated. A very small fraction of variant reads were identified in sequence data from wild-type samples: 0.6% and 0.02 % of total reads in the data for ‘Hawaii 4’ and ‘Calypso’ wild types, respectively. These reads were considered background ‘noise’ as a result of index-switching between Illumina barcodes, an event that occurs at low rates during Illumina sequencing of pooled samples on a single flow cell [50].

In most cases the observed phenotype reflects the sequence data obtained: sequence data for the majority of white samples in both cultivars (23 of 31 samples and 11 of 18 ‘Hawaii 4’ lines; 10 of 14 samples and 4 of 7 ‘Calypso’ lines) lack or have a negligible fraction of wild-type sequence reads (< 0.05%-0.1% of total reads), including some shoot lines for which the fraction of wild-type sequence data for all samples sequenced is negligible (‘Hawaii 4’ lines 12, 14, 79 and 89 and ‘Calypso’ line 1). The exception is ‘Calypso’ shoot 1C, which is probably not transformed (PCR data show absence of a product in reactions using Nos-Kan cassette and sgRNA insert site T-DNA primers. Sequence data for ‘Calypso’ shoot line 7D is anomalous as it was observed to have a variegated phenotype. The presence of a low fraction (<5%) of wild-type reads in data from white phenotypes in some cases may be attributed to subjectivity or difficulty in assessing phenotype (for example where the leaf samples are very small), and may reflect the normal limits of visual assessment for non-functional *PDS* alleles: ‘Hawaii 4’: 10 D1 (very small albino shoot); 28 A1 (white axillary on variegated shoot); 46 F (small albino shoot); 90 B (albino axillary shoot, apical expanded leaf, no green tissue obvious) (Fig. S2), ‘Calypso’: 100 2 (small albino shoot, Fig. S3); 5 Ci (albino leaf from variegated shoot, Fig. 4).

Only one shoot heterozygous for target sequence mutation lacks the expected albino phenotype: sequence data for a green ‘Hawaii 4’ transgenic ‘Hawaii 4’ 3A show that 50% of reads are deletion variants, but indicate that one allele has an amino acid substitution (threonine to asparagine) and that this encodes a functional PDS protein. Published PDS protein sequences [51] for maize and rice also encode asparagine at this position (Fig. S6).

The ranges of wild-type reads commonly observed for pale green and variegated samples are 7-19% (7 samples from 7 lines of ‘Hawaii 4’) and 6-36% (15 samples from 10 lines of ‘Hawaii 4’, 2 samples from 2 lines of ‘Calypso’), which is taken to be due to the proportion of alleles and cells expressing functional PDS protein.

#### 3.2.5 Possible variation in PDS gene allele functionality in Fragaria and gene redundancy in the octoploid genome

In lines of both cultivars, sequence data for some observed phenotypes show unexpected proportions of wild-type and mutated sequences (Table S4, Figs. 4, S2, S3): all ‘Hawaii 4’ transgenic samples with a green phenotype are single-allele sequence variants, with approximately only 20% (‘Hawaii 4’ 46A), and 30% to 60% wild-type reads (‘Hawaii 4’ lines 10 A2, 28 C2, 28 C3, 32 A1 32 A2, 74 A and 90C); also, white phenotypes with higher than expected wild-type sequence fractions (15-30%) include ‘Hawaii 4’ 9B, 28 - A, B, C2, C3, 32 A1, A2, B2, 39 B and 90 (Fig. S2). Nishitani *et al*. [24] observed green transformants in apple (another diploid member of the Rosaceae) with partially mutated sequences of the *PDS* gene. It is possible that one of the *PDS* alleles is silenced or nonfunctional, and the fraction of wild-type sequence reflects the proportion of non-functional *PDS* alleles. ‘Hawaii 4’ samples 36 A3 and 28 D1, which are largely white with green leaf edge sectors and a high fraction of wild-type reads (27% and 14%) are possibly periclinal chimaeras: a mosaic of white and green cells in the L2 layer would be manifested as small green flecks at the leaf edge, and would explain the high wild-type reads in comparison with samples which show small green sectors in the central part of the lamina (e.g. ‘Hawaii 4’ 2 Ci, 32 B1 and 39 C1, with 0.7-3.5% wild-type reads).

The phenotypic and sequence data for ‘Calypso’ suggest that in the octoploid, cultivated strawberry there is only one functional *PDS* gene allele: sequence data for albino shoot 94 includes 31% wild-type reads and a green transgenic shoot (59) has only 11% wild-type reads. Albino or variegated samples with unexpectedly high wild-type reads and complex chimaeric genotypes may include mutations of both functional and nonfunctional alleles (e.g. ‘Hawaii 4’ 28 A1, 46 F, 90 B, and ‘Calypso’ 20 B,100 2-see Supplementary Figures S2 and S3). Evidence for gene silencing and loss of redundant sequences in polyploids is well-documented [52], and gene copy loss and deletion in polyploid *Fragaria* has been described by Rousseau-Gueutin et al.[53]. The fact that a single wild-type amplicon was found is suggestive, but not conclusive that the PDS1 locus is functionally diploid and that complex variants observed in amplicon profiles are due to chimeric regeneration, rather than multiple edits of homeologs.

#### 3.2.6 CRISPR/Cas9 gene edited shoots can be efficiently produced in diploid and octoploid Fragaria

The primary aim of this work was to generate a set of detailed data using a visible marker that will be of relevance for gene editing in *Fragaria* of single- and multi-copy genes for which there is no visible phenotype. We have shown that *via Agrobacterium*-mediated transformation of leaf and petiole explants it is possible to regenerate shoots, which are biallelic mutants for a single copy gene, at a high frequency in both the diploid and octoploid genomes, giving confidence that effective targeting of multiple alleles and gene copies may be achieved. Effective targeting by CRISPR-Cas9 mutagenesis of genes and targeting of multiple homoeologs has been demonstrated in other polyploid crops, such as wheat [28], potato [29] and oilseed rape [30, 31]. Additionally, efficient targeting of multiple gene copies has been shown in rice [32, 34, 35]. In the octoploid strawberry an alternative *U6III* promoter may enhance expression of guide sequences and increase mutagenesis. Zhou et al. [54] have recently reported successful gene editing (of auxin biosynthesis and auxin response factor genes) in wild strawberry, showing similar rates of mutation to those reported here, and demonstrated improved recovery of mutants by use of dual sgRNAs.

In conclusion, when combined with high-throughput screening [55] and high-throughput sequencing, we expect efficient isolation of non-visible mutants of single and multi-copy genes in both diploid and octoploid *Fragaria* to be achievable.

## Author contributions

R.H. and F.W. devised the research plan. F.W. planned the experiments, designed and constructed the CRISPR cassettes and vectors, and performed phenotypic and molecular analysis of transgenic lines. K.H. produced transgenic plants. A.D.A. performed bioinformatic analysis of sequence data and generated variant allele summary plots. F.W. wrote the manuscript with input from R.J.H., A.D.A. and A.J.S.

## Acknowledgements

This work was supported by the Biotechnology and Biological Sciences Research Council (BBSRC) (BB/K017071/2 and BB/N006682/2). The authors thank Maria Sobczyk for assistance with obtaining the *PDS* gene sequences for *F. vesca* and *F*. x *ananassa*, and Helen J Bates for assistance with Illumina sequencing. The binary vector pFGC-pcoCas9 was a gift from Jen Sheen (Addgene plasmid # 52256). pGreen plant selection and marker cassettes were provided by Mark Smedley, John Innes Centre, Norwich, Norfolk, United Kingdom.

